# Effect of rapamycin on mitochondria and lysosomes in fibroblasts from patients with mtDNA mutations

**DOI:** 10.1101/2021.03.31.437507

**Authors:** Nashwa Cheema, Jessie M. Cameron, David A. Hood

**Author notes:** **To whom correspondence should be addressed:** David A. Hood, PhD, Muscle Health Research Centre, School of Kinesiology and Health Science, York University, 4700 Keele St., Toronto, ON, M3J 1P3, Canada.

## Abstract

Maintaining mitochondrial function and dynamics is crucial for cellular health. In muscle, defects in mitochondria result in severe myopathies where accumulation of damaged mitochondria causes deterioration and dysfunction. Importantly, understanding the role of mitochondria in disease is a necessity to determine future therapeutics. One of the most common myopathies is **m**itochondrial **e**ncephalopathy **l**actic **a**cidosis **s**troke-like episodes (MELAS), which has no current treatment. Recently, MELAS patients treated with rapamycin exhibited improved clinical outcomes. However, the cellular mechanisms of rapamycin effects in MELAS patients are currently unknown. In this study, we used cultured skin fibroblasts as a window into the mitochondrial dysfunction evident in MELAS cells, as well as to study the mechanisms of rapamycin action, compared to control, healthy individuals. We observed that mitochondria from patients were fragmented, had a 3-fold decline in the average speed of motility, a 2-fold reduced mitochondrial membrane potential and a 1.5–2-fold decline in basal respiration. Despite the reduction in mitochondrial function, mitochondrial import protein Tim23 was elevated in patient cell lines. MELAS fibroblasts had increased MnSOD, p62 and lysosomal function when compared to healthy controls. Treatment of MELAS fibroblasts with rapamycin for 24 hrs resulted in increased mitochondrial respiration compared to control cells, a higher lysosome content, and a greater localization of mitochondria to lysosomes. Our studies suggest that rapamycin has the potential to improve cellular health even in the presence of mtDNA defects, primarily via an increase in lysosomal content.

## Introduction

Rapamycin, an autophagy inducer and a pharmacological inhibitor of mammalian target of rapamycin (mTOR), extends the lifespan of mice and delays the onset of age-associated diseases in model organisms by rejuvenating cellular health (1). Treatment with rapamycin has ameliorated adverse phenotypes and improved endurance exercise and mitochondrial function. The beneficial effects are due to an increase in autophagic flux, an upregulation of lysosomal biogenesis, a higher abundance of Lamp1 protein and the localization of TFEB, the master regulator of lysosomal biogenesis, to the nucleus (2–4). Lysosomes play an integral role in maintaining mitochondrial health by degrading damaged mitochondria in the process of mitophagy. Recent studies have suggested that mTOR has an additional role in mitophagy(5), wherein global autophagy induction can trigger the activation of mitophagic pathways in the presence of dysfunctional mitochondria (6, 7).

Mitochondrial myopathies are a group of neuromuscular disorders with prominent organelle dysfunction due to genetic abnormalities. Symptoms exhibited by myopathy patients include muscle weakness, exercise intolerance, mobility impairments, heart failure, stroke-like episodes, seizures, deafness, blindness and dementia. Common mitochondrial myopathies include Kearns-Sayre syndrome (KSS), myoclonus epilepsy with ragged-red fibers (MERRF), and mitochondrial encephalomyopathy with lactic acidosis and stroke-like episodes (MELAS). MELAS is one of the most common maternally-inherited mitochondrial disorder, with accumulations of point mutations in the mtDNA (8, 9) and a prevalence rate estimated to be 6 per 10,000 people (10). As mitochondria are essential to nearly every cell in the body, the disease affects the brain, heart, muscles, gastrointestinal tract, endocrine function, pulmonary and renal systems (11, 12). Due to such a wide range of symptoms and genetic features, conducting controlled clinical trials with therapeutic agents is challenging (12–16). No treatments have been consistently associated with an improvement in clinical outcomes (14). Hence, the lack of a uniform treatment strategy provides a strong need for research in a potential therapeutic for MELAS.

Until recently, rapamycin was only widely used in genetic and mammalian models of disease. Currently however, multiple clinical trials are now underway to test the efficacy of rapamycin in various human diseases (17, 18). Treatment of patients with MELAS resulted in improved health progression, and rapamycin rescued some mitochondrial defects in isolated cultured fibroblasts (17). However, the cellular mechanisms of how rapamycin acts to improve cellular health in the presence of mtDNA defects are currently unknown. Thus, in this study we sought to investigate the extent of mitochondrial dysfunction in fibroblasts isolated from MELAS patients, as well as the effect of rapamycin treatment on mitochondria and lysosomes, since the function of these organelles is inextricably tied (19).

## Methods

### Cell culture

Human skin fibroblasts from the forearm were obtained with informed consent from family members for diagnostic purposes, with approved methodologies established at the Hospital for Sick Children, Toronto, Canada. Cells cultures were established from 4 patients as well as 3 controls and were maintained as frozen stocks until used for analyses. The genetic mutation and patient demographics are reported in Table 1. The cells were thawed and grown at 5% CO_2_ at 37° C in AMEM supplemented with 10% Fetal Bovine Serum and 1% Penicillin/ Streptomycin until they reached 80-90% confluency, as done previously (20, 21). Cells were treated with 100 nM rapamycin (Life Technologies, Burlington, ON, CA) or vehicle (DMSO) for 24hrs. To visualize mitochondria, fibroblasts were transfected with the plasmid mito-dsRED2, red fluorescent protein fused with the mitochondrial targeting sequence from cytochrome c oxidase subunit VIII (Clontech, Mountain View, CA) using Lipofectamine 2000 (Life Technologies, Burlington, ON, CA).

**Table 1.** Age, sex and mutation identified from forearm samples in four MELAS patients.

|  | Patient ID<br>Number | Age | Sex | Mutation | Heteroplasmy |
| --- | --- | --- | --- | --- | --- |
| P1 | 5509 | 6 months | M | G13513A | 70% |
| P2 | 13801 | >18 years | F | T8993C | 72 or 82% |
| P3 | 21124 | 2 years | F | T8993C | >95% |
| P4 | 20551 | 2 years | F | G13513A | unknown |

|  | **Patient ID**<br>**Number** | **Age** | **Sex** | **Mutation** | **Heteroplasmy** |
| --- | --- | --- | --- | --- | --- |
| P1 | 5509 | 6 months | M | G13513A | 70% |
| P2 | 13801 | >18 years | F | T8993C | 72 or 82% |
| P3 | 21124 | 2 years | F | T8993C | >95% |
| P4 | 20551 | 2 years | F | G13513A | unknown |

### Live cell imaging

To visualize nuclei, mitochondria and lysosomes, fibroblasts were incubated with Hoechst 33342 (ThermoFischer, MA, USA), 10 nM mitotracker green (Invitrogen, MA, USA) and 10 nM lysotracker red (Invitrogen, MA, USA) for 15 mins at 5% CO_2_ at 37° C. Stained and mtDsRed transfected cells were visualized using an inverted Nikon Eclipse TE-2000 confocal microscope equipped with a ×60/1.5 oil objective lens and a custom-designed chamber for live cell imaging that maintained a constant temperature of 37°C with 5% CO_2_. Live cell time lapse imaging of mitochondrial dynamics was captured at 2 second intervals for a total time of 5 mins, as done previously (22).

### Analysis of mitochondrial motility and morphology

Analyses were performed using NIS Element AR 3.1 software (Nikon Inc., USA). Mitochondrial motility was calculated by kymographs for each fibroblast, using 7-11 cells per cell line. The average motility for each fibroblast is reported by the measuring speed of 8-15 mitochondria per cell. Kymographs were generated from peripheral regions of each cell as individual mitochondria could be resolved for analysis. Mitochondria in tubular networks, often surrounding the nucleus, were not included for analysis. Velocity measurements were exported from NIS software. Calculations of the percent fused and fragmented mitochondria were made using the General Analysis program in NIS Elements where the minimum size of mitochondria was set at 0.2 μm.

### Lysosomal number and colocalization coefficient

Both lysosomal and nuclear number were determined from confocal images from each cell line (10/20x images) and analyzed in ImageJ (NIH, USA) via the count tool. Lysosomal number was normalized to the number of nuclei. Pearson coefficient of lysotracker red and mitotracker green stained fibroblasts was obtained from NIS Elements. For each cell line, one coverslip was stained, and three images were taken at different locations of the coverslip. For each image, the colocalization coefficient value was obtained (n=3 for each cell line). Vehicle-DMSO and rapamycin treated cells were imaged on the same day.

### Lysosomal function

To determine lysosomal function, fibroblasts were incubated with a fluorogenic substrate for proteases, DQ Red BSA (Invitrogen, MA, USA). The dye conjugated to BSA is non-fluorescent when the protein is in its native state. When cleaved by proteases in the lysosome, the fluorescent quenching effect is relieved and single peptides emit red fluorescence. Fluorescence was quantitated by flow cytometry, as described below.

### Flow Cytometry

The fluorescent geomean intensities of mitotracker green, lysotracker red, DQ Red BSA and JC-1 were determined using a BD FACSCaliber flow cytometer (BD Biosciences, CA, USA) equipped with FL-1 (green), FL-2 (red) and FL-3 (deeper red) detectors.

### JC-1 Fluorescence

Fibroblasts were grown to 80% confluency. Cells were washed with PBS and incubated with 1.5 μM of JC-1 at 37 °C in the dark for 30 mins. After incubation, cells were washed with warm PBS and harvested. Pellets were resuspended in phenol red-free media and analyzed in the flow cytometer.

### DCF Fluorescence

Fibroblasts were seeded at 20,000 cells in a 96-well optical bottom plate. After 24 hrs, cells were incubated with 2’7’ dichlorofluorescin diacetate (DCF) (Invitrogen, USA) at 37°C for 30 min. Fluorescence (excitation 480nm, emission 520 nm) was measured using a Synergy HT microplate reader (BioTek Instruments,VT, USA). After incubation, cells were washed with PBS and incubated with DAPI stain at 37°C for 15 min. Fluorescence was measured using the microplate reader and data were normalized to DAPI staining per well.

### Mitochondrial Respiration

Assays were performed using a Seahorse XF96 Cell Mito Stress Test Kit (Agilent Biosciences, USA). Briefly, fibroblasts were seeded at a density of 20,000 cells per well in a 96-well Seahorse cell culture plate and incubated overnight. Each fibroblast cell line was seeded in 12 replicate wells. In experiments involving drug treatments, 6 wells per cell line were treated with DMSO or rapamycin. After 24h, the Seahorse XF96 Extracellular Flux Analyzer was used to measure the oxygen consumption rate (OCR) of each well. After the OCR was measured, the seeded plate was washed twice with PBS. Cell lysates were extracted from each well and Bradford reagent was added to measure protein absorbance per well. The assays were analyzed using XF Wave software and all OCR measurements were normalized to protein absorbance.

### Western blotting

Cell protein extracts were prepared and separated in polyacrylamide SDS-PAGE gels. Proteins were then transferred onto nitrocellulose membranes and blocked in 5%-10% milk in TBS-T. Subsequently, membranes were incubated with the appropriate concentration of primary antibodies overnight at 4°C, and then with HRP-conjugated secondary anti-mouse (Cell Signaling 7076S) or anti-rabbit antibodies (Cell Signaling 7074S) for 1hr at room temperature. Membranes were then visualized with enhanced chemiluminesence using a Carestream Imaging system. Primary antibodies used were anti-Tim23 (BD Biosciences BD611222), anti-α smooth muscle Actin (Abcam ab5694), anti-p62 (Abcam ab56416), anti-B1/2 V-ATPase (SantaCruz SC-55544), anti-Kif5B (SantaCruz SC 28538), or anti-MnSOD (Upstate 06-984). All antibodies were diluted 1:1000. The blots were cut into four parts. The loading control was obtained from the same blot for all the antibodies. The p62 immunoblot was reprobed for V-ATPase and the Tim23 immunoblot was stripped and reprobed for MnSOD, which migrate at different molecular weights.

### Statistical analysis

Data are presented as means +/- SD for all control and patient fibroblast lines. All statistical analysis was performed using GRAPHPAD PRISM version 9.00 for Windows (GraphPad Software, San Diego, CA, USA). Unpaired t-tests comparing average control to average patient were performed in Figures 1–3. In Figure 4, a paired t-test (denoted as #) was used for analysis when comparing vehicle vs rapamycin treatment of respective cell lines. An unpaired t-test, denoted as *, was used for comparison of average vehicle vs average rapamycin treatment. In Figure 5, a one-way ANOVA with a Tukey post-hoc test was performed for analyses. In all statistical analyses, a p-value < 0.05 was considered significant.

**Figure 1.**
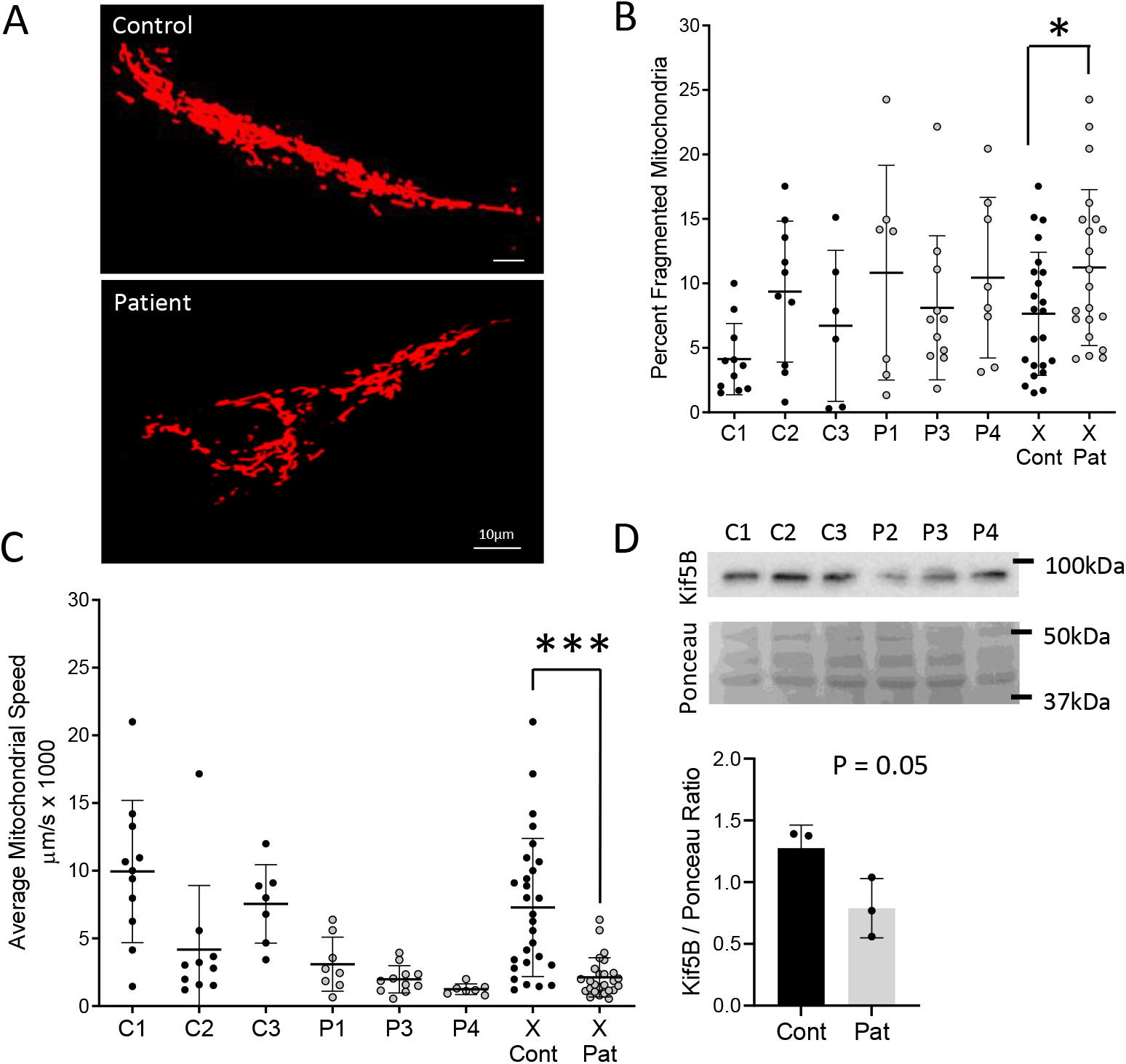
Mitochondrial motility and morphology in fibroblasts isolated from human control individuals and MELAS patients. A) Mitochondrial network visualized by mtDsRed in control and patient representative fibroblast. Live cell imaging was performed with a confocal microscope equipped with a heated stage and CO_2_ chamber. B) Percent of fragmented mitochondria in cells from each cell line, control and patient (n=7-12 cells per cell line for control and patient). All cell lines were included in the average control (Cont) and patient (Pat). Average percent of fragmentation is shown for all control/patient cell line. Control represented by black bar and patient cell lines by grey bar. Motility of mitochondria was measured via kymographs to determine (C) average speed. Kymographs were generated from time lapse videos of 2 sec intervals for 5mins duration. N= 6-12 cells per cell line and 8-15 mitochondria per cell. The average control had n=24 and average patient n=21 cells. D) Kif-5B protein level in three control and three patient cell lines. Unpaired t-test was performed where * p-value < 0.05 and *** p-value is <0.0001.

## Results

### Mitochondria are fragmented and motility is reduced in MELAS fibroblasts

To assess the morphological integrity of mitochondria in fibroblasts isolated from healthy individuals and MELAS patients, cells were transfected with mtDsRed, a fluorescent mitochondrially-targeted protein, and visualized by confocal microscopy. A representative cell from control and MELAS fibroblast lines is shown in Figure 1A, illustrating the degree of mitochondrial network fragmentation in patient cells. A higher percent of fragmentation was observed in all three MELAS fibroblast lines when compared to the controls, with an average 1.5-fold increase in MELAS fibroblasts (Fig. 1B). To determine whether this fragmentation had an impact on mitochondrial motility, 5-min time lapse videos were generated by live cell imaging of mtDsRed transfected cells. Motility was calculated by generating kymographs for each fibroblast. The average motility is reported by measuring the speed of 8-15 mitochondria for each fibroblast. All MELAS fibroblasts lines had reduced average (Fig. 1C) mitochondrial speed, with a reduction greater than 2-fold when compared to the control fibroblast lines. To assess whether this effect could be due to the expression of motor proteins responsible for organelle motility along microtubules, we measured the level of the kinesin motor protein isoform Kif5B by western blots in control and patient fibroblasts. A decline (p=0.05) was observed in patient cell lines (Fig. 1D), which could contribute to this reduction in motility.

### MELAS fibroblasts exhibit reductions in mitochondrial membrane potential and respiration, but no change in mitochondrial content

To verify changes in mitochondrial function in MELAS cells, we first determined the mitochondrial membrane potential of control and MELAS fibroblast lines. JC-1 is a fluorescent probe that remains in the cytosol as a monomer and emits green fluorescence when mitochondria have lost their membrane potential. The dye aggregates within mitochondria to emit red fluorescence where there is an intact membrane potential. Thus, the ratio of red to green fluorescence is an indicator of mitochondrial health within the cell. MELAS fibroblast lines exhibited a significant decline in the JC-1 ratio, indicating the loss of mitochondrial membrane potential in comparison to the control cells (Fig. 2A). To assess mitochondrial respiration, we measured the oxygen consumption rate (OCR) of our fibroblast lines using Seahorse. A typical OCR profile, along with the drug additions and resulting data is shown in Fig. 2B. There was a significant decline in basal (Fig. 2C) and proton linked (Fig. 2D) mitochondrial respiration in all MELAS cell lines. Maximal respiration, which is an indication of total mitochondria in the cell, did not differ between control and MELAS cells (Fig. 2C). Patients displayed a significantly increased Tim23 protein expression relative to controls (Fig 2E) suggesting a compensatory adaptation in the import pathway, to correct for the mitochondrial dysfunction observed.

**Figure 2.**
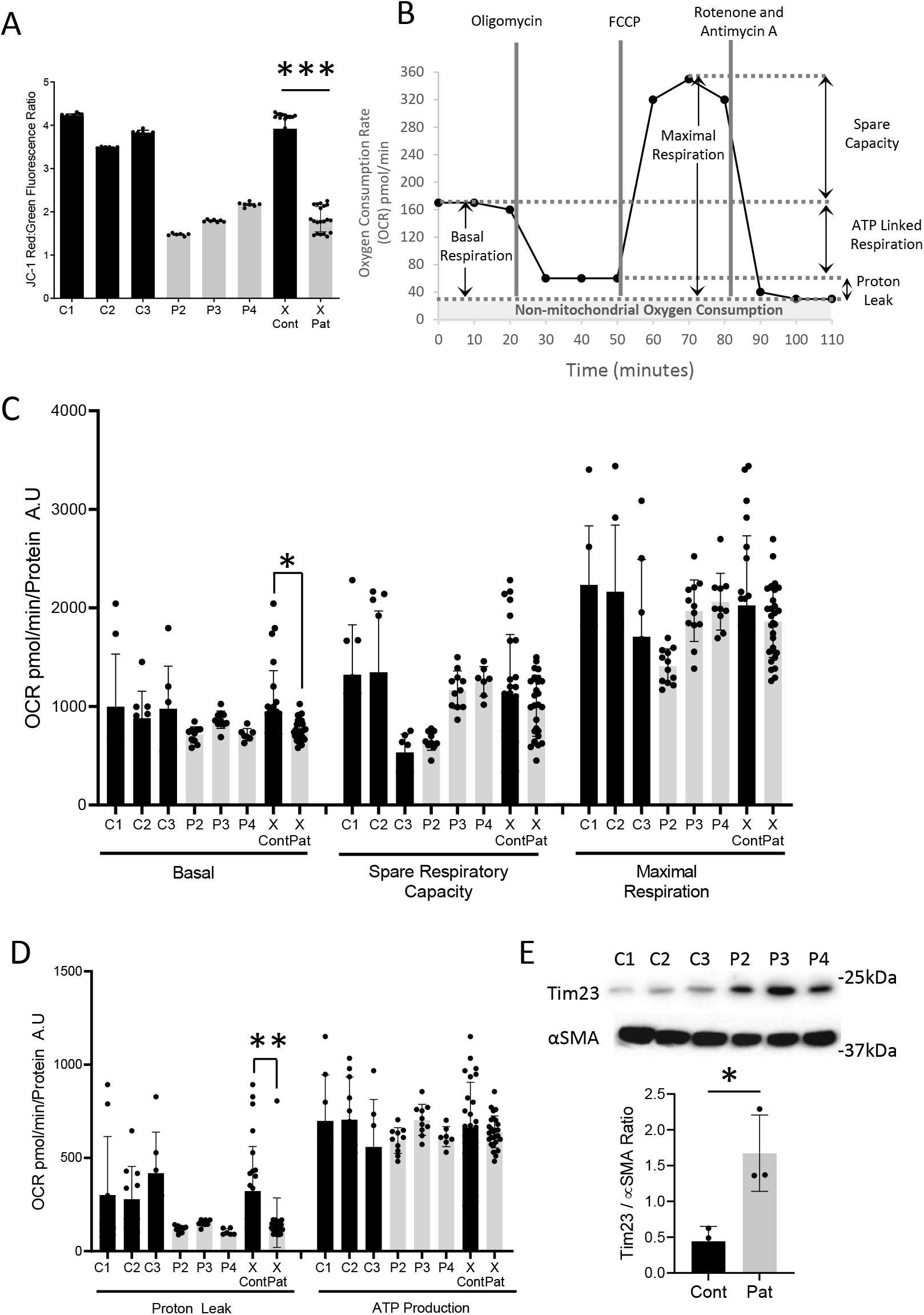
Mitochondrial function and amount in healthy and MELAS fibroblasts. A) Mitochondrial membrane potential as measured by fluorescent probe JC-1. Aggregated form of JC-1 emits a red fluorescence in mitochondria with an intact membrane potential whereas a monomer JC-1 fluoresces green when present in the cytosol. Fluorescent intensities were measured in the flow cytometer for control (Cont) and patient (Pat). B) A diagram of an annotated seahorse XF Analyzer graph with different measured parameters labeled for Mitostress assay C) Average basal, spare respiratory capacity and maximal respiration in three control and three MELAS fibroblast cell lines, n=7. D) Average Proton leak dependant respiration and ATP Production in three control and three MELAS fibroblast cell lines, n=7. E) Tim23 levels three control and three patient cell lines. Smooth muscle actin, α-SMA, used as loading control. Unpaired t-test was performed where * p-value < 0.05, ** p-value < 0.01 and *** p-value is <0.0001.

### MELAS fibroblasts have increased protease function and oxidative stress

Lysosomes play an integral part in maintaining cellular health by degrading dysfunctional organelles, like mitochondria. We determined lysosomal content in fibroblasts using flow cytometry following incubation with the fluorescent dye, lysotracker red. No differences were detected between control and MELAS fibroblast cell lines (Fig. 3A). This corresponded to the lack of difference observed between control and patients in the lysosomal protein marker B1/2 V-ATPase (Fig. 3C, D). Lysosomal function of live cells was also assessed using the fluorescence intensity of a proteolytically cleaved fragment generated in the lysosome (Fig. 3B). Despite individual differences, two out of the three MELAS cell lines exhibited clearly elevated lysosomal protease function compared to control fibroblasts (Fig. 3B). In order to match this with estimates of autophagy, p62 was used as a surrogate indicator of autophagy flux in these cells. The content of p62 was elevated in the patients by about 30% (p<0.05; Fig. 3C, D), suggesting that while lysosomal function was enhanced, uptake or delivery of autophagic substrates into the lysosome could be impaired in patient cells. Since ROS can act as a signal for the activation of autophagy, we used DCF fluorescence as an indicator of ROS emission, and MnSOD protein as a measure of anti-oxidant capacity in these cells. Our data indicate that these patient cells exhibited a marked 7-8-fold increase (P<0.05; Fig. 3E) in DCF fluorescence compared to controls, an increase similar to previous reports (21), which was only accompanied by a 2-fold higher level (P<0.05; Fig. 3D) of MnSOD protein.

**Figure 3.**
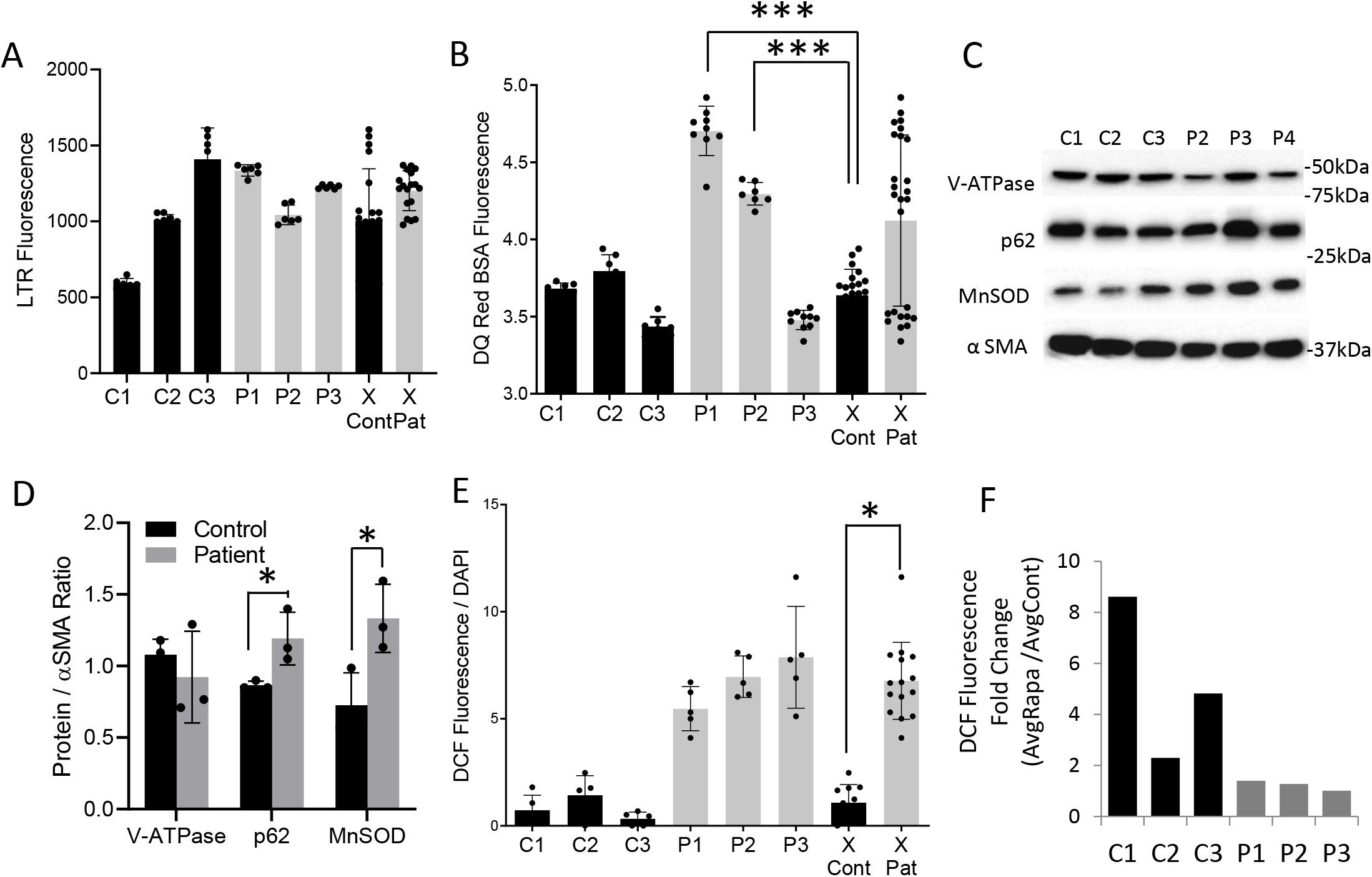
Lysosomal function and oxidative stress in healthy and MELAS fibroblasts. A) Fluorescent intensity for lysotracker red and B) lysosomal function via DQ Red BSA in control and patient fibroblasts, n=7. All cell lines were included in the average control (Cont) and patient (Pat). No significance (n.s) when comparing lysosomal function between average control and patient fibroblasts. C) B1/2 v-ATPase, p62, MnSOD protein expression in three control and three patient cell lines. Smooth muscle actin, α-SMA, used as loading control. D) Normalized protein expression in control and patient cell lines (n=3). E) ROS measured with DCF fluorescent probe in control and patients. F) Fold change of ROS levels upon rapamycin treatment. n=5 for each cell line treated with DMSO or rapamycin. Unpaired t-test was performed where * p-value < 0.05 and *** p-value is <0.0001 patient vs control.

### Rapamycin rescues basal and proton leak mitochondrial respiration in MELAS fibroblast

We then assessed mitochondrial function following treatment of the cells with rapamycin for 24 hours. Upon addition of rapamycin, control fibroblasts displayed 2-9-fold increases in DCF fluorescence. However, DCF fluorescence in MELAS fibroblasts did not increase in response to rapamycin (Fig. 3F). In addition, while rapamycin treatment had very little effect on OCR parameters measured in control cells, apart from a minor reduction in basal respiration (Fig. 4A, B), rapamycin treatment significantly improved the respiratory function of MELAS fibroblasts, by increasing basal (60%, P<0.05), as well as proton leak (100%, P<0.05; Fig. 4C,D) respiration.

**Figure 4.**
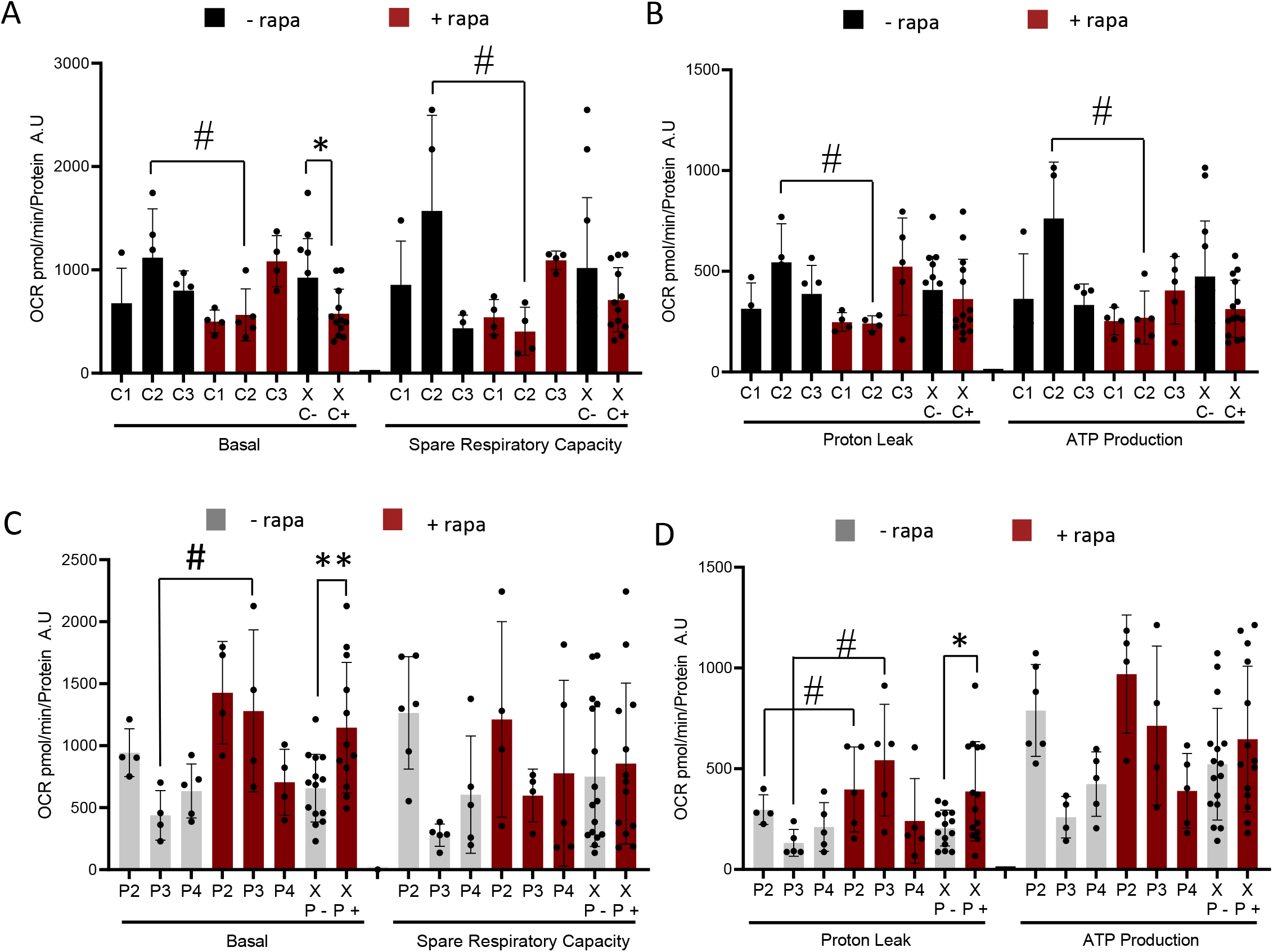
Rapamycin effects mitochondrial respiration in fibroblasts. A-B) Average basal respiration, spare respiratory capacity, proton leak and ATP production for all control fibroblast lines with and without rapamycin treatment. C-D) Basal, spare respiratory capacity, proton leak and ATP production for all MELAS lines with and without rapamycin, n= 4-6 for each cell line treated with DMSO or rapamycin. All cell lines were included in the average control (C - / + rapamycin) or patient (P - / + rapamycin) for a n=11. Paired t-test was performed where # p-value < 0.05 rapamycin treatment (+rapa) vs vehicle-DMSO treatment (-rapa) of respective cell line. Unpaired t-test was performed where * p-value < 0.05 and ** p-value <0.01 of average vehicle vs average rapamycin treatment.

### The beneficial effect of rapamycin on mitochondria could be mediated by increased lysosomal number

It has been known for some time that rapamycin treatment leads to an elevation in lysosome content (2–4). To evaluate this in patient and control cells, we stained fibroblasts with lysosomal and nuclear markers (Fig. 5A). We observed that rapamycin induced a 4-5-fold increase (P<0.05) in lysosomal number in both control and patient cells when compared to vehicle treatment (Figs. 5B). Rapamycin had no significant effect on lysosomal function in either control or patient cell lines (Fig. 5C). Our data also show that, upon rapamycin treatment, lysosomes colocalize to a much greater extent with mitochondria in MELAS fibroblasts (Fig. 5D). This increase in colocalization was not evident in control cells. These data suggest that the rapamycin-induced increase in lysosome content may help traffic dysfunctional mitochondria to the organelle for clearance. This would serve to reduce the pool of dysfunctional organelles and improve mitochondrial function in MELAS patient cells, as evident from better respiratory rates and a lack of increase in ROS emission, when treated with the drug.

**Figure 5.**
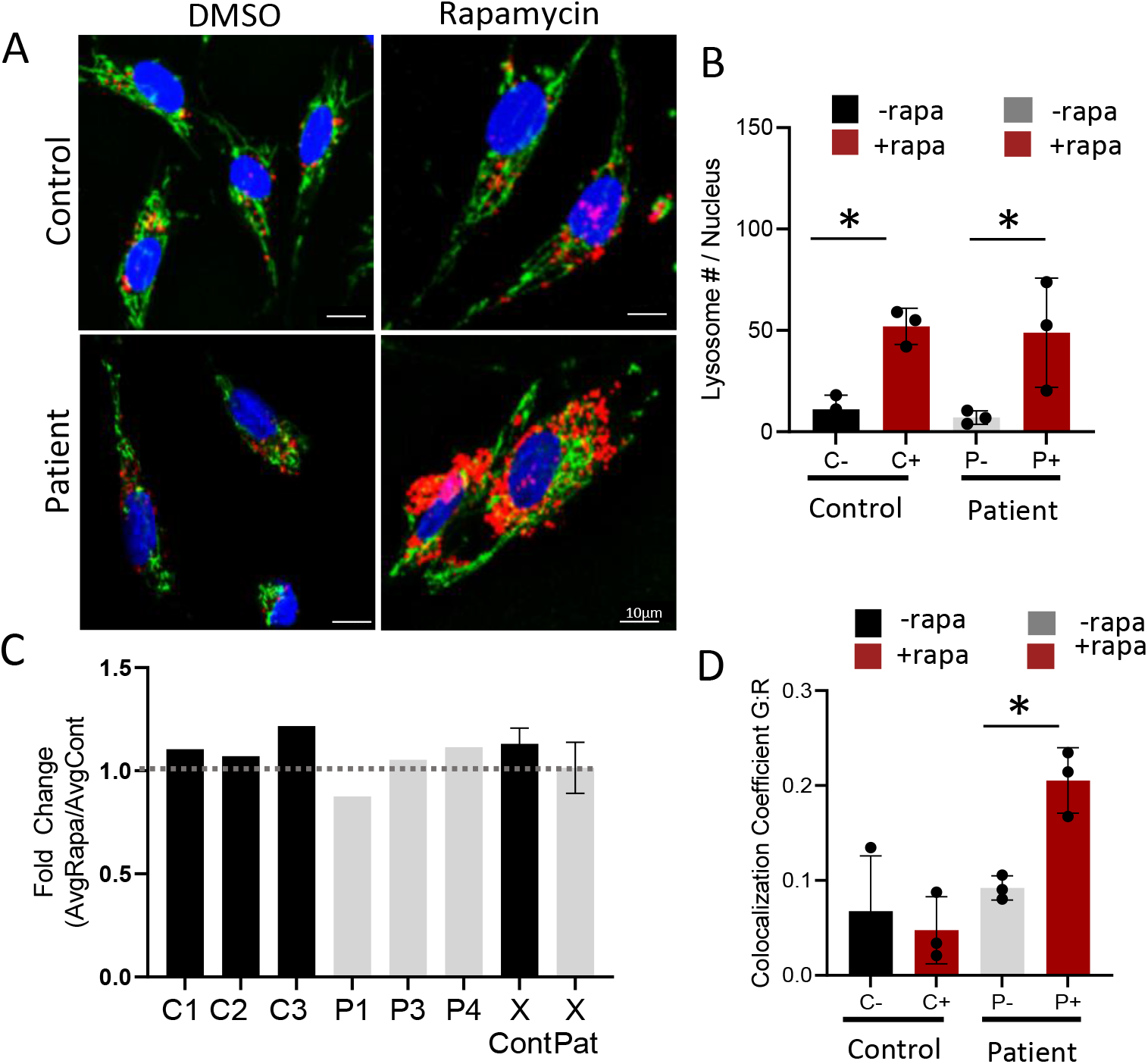
Rapamycin effect mediated through lysosomal alterations. A) Representative confocal image of hoechst (H), mitotracker green (MTG) and lysotracker red (LTR) staining from one control and patient treated with DMSO or rapamycin. B) Quantification of lysosomal number from DMSO and rapamycin treatment in control and patient cell lines (n=3). Rapamycin treatment increased lysosomal number per nucleus in control fibroblast lines. C) Rapamycin effect on lysosomal function in control and MELAS fibroblast lines, n=7. All cell lines were included in the average control (Cont) and patient (Pat). D) LTR signal colocalizes with MTG with a higher colocalization coefficient in MELAS lines treated with rapamycin. Images were taken from different locations on a coverslip, n=3 images for each cell lines. 1-way ANOVA was performed where * p-value < 0.05 of average rapamycin treated vs average vehicle-DMSO treatment.

## Discussion

Mitochondrial defects can cause severe energy deficits and cell death. To preserve health, the maintenance of mitochondrial integrity and function is critical. Potential therapeutic strategies for mitochondrial disorders are being investigated, however there is currently no effective intervention. Rapamycin has shown benefits for muscle myopathies (2, 17, 23) but the mechanism of action is not fully known. Our data suggest that as an autophagy inducer, rapamycin may improve mitochondrial dysfunction, since we observed that mitochondrial respiration in MELAS cells was, in part, rescued by rapamycin treatment.

Mitochondrial dysfunction is a common feature of mitochondrial myopathies such as MELAS (23–25). We observed a ~50% reduction in membrane potential in patient cells, similar to other reported literature in MELAS fibroblasts (23). Along with lower mitochondrial membrane potential, mitochondrial respiration was reduced with no detectable changes in mitochondrial content (26). Another recent study (27) has reported no changes in ATP production in MELAS fibroblasts. Similarly, we detected no differences in ATP-linked respiration in our cell lines. An indirect measure of mitochondrial content in a cell is maximal respiratory capacity, which is not different between our control and patient fibroblasts. Studies have generally reported no changes in total mitochondrial mass in MELAS fibroblasts (21, 26) and have observed an upregulation of mitochondrial import machinery (20). We noted an increase in Tim23, a mitochondrial protein encoded by the nuclear genome, similar to previous studies in mtDNA disease patients (20). Higher levels of Tim23 might be an indication of a compensatory retrograde signaling pathway activated in the presence of mitochondrial dysfunction. In the analysis of multiple MELAS patients, individual differences in pathophysiology have been observed, whereby some patients exhibit an extreme phenotype of reduced mitochondrial potential, respiration and mitochondrial proteins, whereas others have less drastic changes, with no differences detected (23). This is likely due to variability in patient symptoms, age of onset and manifestations, and this could clearly account for some of the variability observed between patient cells in the current study. Other limitations that could have introduced variability among patient and control cells include different passage numbers, as well as cell proliferation rates.

Mitochondrial motility is necessary to maintain a healthy pool of the organelle. Cytoskeletal and motor proteins allow for the transport of mitochondria to diverse cellular regions in an effort to maintain reticular, network-like structures. We have previously shown that situations of high oxidative stress led to mitochondria becoming stationery and fragmented (22). Reduced mitochondrial motility has been reported in a variety of metabolic, neurodegenerative and mitochondrial diseases (28). Our data indicate that MELAS fibroblasts exhibit fragmented mitochondria, as reported by others (23), and have a reduced average motility. This is likely due in part to lower levels of the kinesin isoform Kif5B, as well as elevated antioxidant enzymes, suggestive of increased oxidative stress. Fragmented, dysfunctional mitochondria are ideal substrates for the mitophagy pathway, leading to their subsequent degradation within lysosomes.

Earlier studies using electron microscopy and flow cytometry have documented an increase in lysosomal number in MELAS cells (26). More recently, investigations have identified the localization of mitochondria to autophagosomes, suggesting mitochondrial degradation (23). While we did not observe an increase in lysosomal content, either visualized using lysotracker staining, or a well-established marker of lysosome content, B1/2 V-ATPase, our data do indicate that protease function within lysosomes was elevated in MELAS fibroblasts. This would suggest an enhanced capacity for cargo degradation in MELAS cells. However, p62 levels were increased in the patients, and this is most often interpreted as an indicator of reduced autophagy flux. This diminished autophagy is supported by the literature which shows that patient cells have elevated levels of autophagy markers such as ATG12-ATG5 and LC3, as well as an accumulation of autophagosomes (23, 24).

The mammalian target of rapamycin mTOR has a well-established role in regulating autophagy. This may also extend to mitophagy (5). The activation of autophagy using rapamycin-induced mTOR inhibition can trigger the activation of mitophagic pathways when there are dysfunctional mitochondria, in a dose- and duration-dependent manner (6, 7). Chronic treatment with rapamycin improved disease phenotype of cells from patients with Leber’s hereditary optic neuropathy (LHON), a mitochondrial disorder. Rapamycin treatment induced autophagy as detected by autophagosome and mitochondrial specific markers co-localization. Furthermore, the threshold of pathogenic mtDNA mutations decreased, suggesting that mitophagy selectively degraded dysfunctional mitochondria (6). We observed a similar increase in lysosomal number in control and patient cell lines with rapamycin treatment. However, in patient cells, lysosomes appeared to be larger in size, had increased staining intensity, and the index of colocalization with mitochondria was significantly elevated in patient cells as a result of rapamycin treatment. This increased colocalization may have resulted in mitochondrial repair, as we observed an increase in organelle function upon rapamycin treatment, but only in patient fibroblasts. This was evident from some improvements in mitochondrial respiration. In addition, notwithstanding the limitations of using DCF as an indicator of ROS (17), the lack of increase in DCF fluorescence in response to rapamycin suggests the induction of a more efficient electron transport chain (29) and a healthier antioxidant status within the MELAS cells (30).

In addition to studies involving cells in culture, the benefits of rapamycin treatment have been observed in *ex vivo* and *in vivo* models. In mice exhibiting extreme myopathy, rapamycin treatment improved the adverse phenotype. Treated mice exhibited a reversal of mitochondrial structural abnormalities, mitochondrial dysfunction and an improvement in muscle function. Rapamycin increased autophagic flux in mice and induced TFEB translocation to the nucleus, which results in lysosomal biogenesis (2). In a recent clinical trial, MELAS patients who were switched to rapamycin treatment exhibited improved metabolic function with less oxidative stress. Furthermore, rapamycin activated autophagy and attenuated the decline in mitochondrial membrane potential and fragmentation of mitochondria (17).

The patients’ cells in the current study have the same mitochondrial content as control cells, but possess mitochondria that are fragmented and poorly motile, with indices of reduced membrane potential, higher ROS emission which outstrips antioxidant capacity, and lower basal respiration. These mitochondria should be recycled and removed to enhance the quality of the organelle pool. Clearance would be achieved with an enhanced rate of mitophagy directed to the lysosome. In MELAS patients, lysosome number is similar as in control cells, but these lysosomes have modestly enhanced lysosome proteolytic function, which seems insufficient to reduce mitochondrial dysfunction. In addition, the lack of mitochondrial motility, perhaps as a result of reduced cytoskeletal transport and enhanced oxidative stress (22), impairs mitochondrial trafficking to the lysosome. Evidence for impaired delivery is found in the increased level of p62, an autophagy substrate. Attainment of a healthier mitochondrial pool could be achieved by providing the cell with a greater number of lysosomes, thereby reducing the traffic jam. Rapamycin treatment clearly increased lysosome content, providing a significantly improved destination for the removal of impaired mitochondria, evident by the co-localization index. This greater lysosomal degradation of dysfunctional mitochondria likely contributes to the better respiration rates back toward healthy control cells, an enhancement that might be amplified with a longer rapamycin treatment time. We speculate (Fig. 6) that the maintenance of a constant mitochondrial content between patient and control cells also obligates a higher degree of retrograde signaling in the presence of rapamycin from mitochondria to the nucleus in patient cells, directing gene expression toward enhanced biogenesis in the face of increased mitochondrial degradation in lysosomes. This is evident in non-treated cells by the higher expression of Tim23, and has been documented previously (20) in MELAS cells. Thus, in summary, our pre-clinical data suggest that rapamycin may have some benefits for MELAS patients with the potential to rescue mitochondrial deficits, primarily via enhanced lysosomal content.

**Figure 6.**
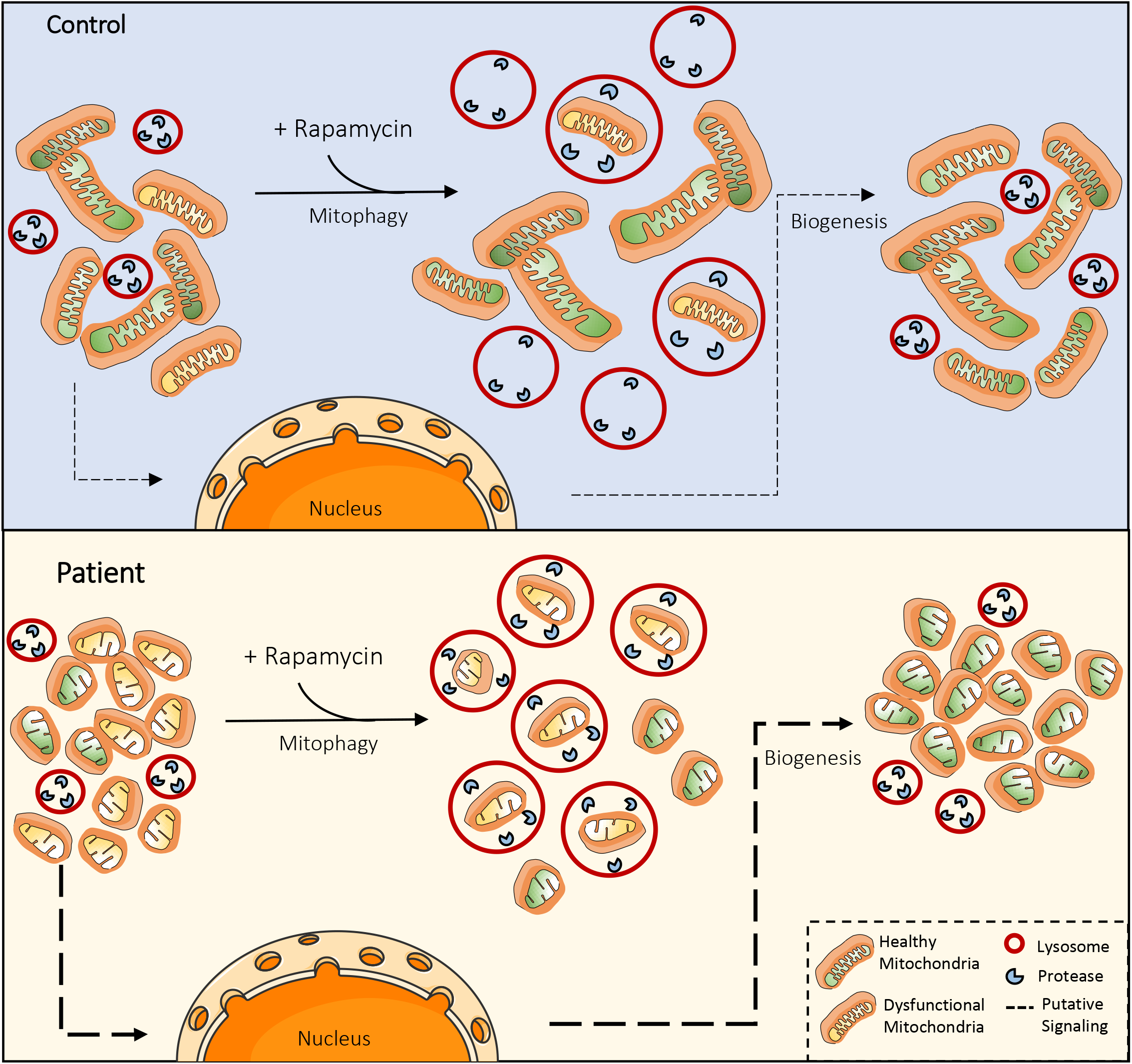
Illustration of the maintenance of mitochondrial health in control and patients with mtDNA mutations. In comparison to healthy controls, mtDNA patients exhibit a similar total mitochondrial content, but these organelles are fragmented with elevated ROS emission, lower membrane potential and basal respiration, and poor cellular motility. To compensate for loss of mitochondrial function, patients have an increased mitochondrial retrograde signaling the nucleus, altering gene expression, as shown previously (20). Lysosome content is similar in control and patient cells, and the addition of rapamycin increases lysosomes in both cell types equally. This effect improves mitochondrial quality in patient cells, promoting mitophagy as evident from greater colocalization of lysosomes and mitochondria, and improved organelle function.

## Acknowledgments

This research was funded by NSERC Canada. David A. Hood is the holder of a Canada Research Chair in Cell Physiology.

## References

1. Kennedy BK, and Lamming DW. The Mechanistic Target of Rapamycin: The Grand ConducTOR of Metabolism and Aging. Cell Metabolism 23: 990–1003, 2016.

2. Civiletto G, Dogan SA, Cerutti R, Fagiolari G, Moggio M, Lamperti C, Benincá C, Viscomi C, and Zeviani M. Rapamycin rescues mitochondrial myopathy via coordinated activation of autophagy and lysosomal biogenesis. EMBO Molecular Medicine 10: e8799, 2018.

3. Sato M, Seki T, Konno A, Hirai H, Kurauchi Y, Hisatsune A, and Katsuki H. Rapamycin activates mammalian microautophagy. Journal of Pharmacological Sciences 140: 201–204, 2019.

4. Zhang X, Chen W, Gao Q, Yang J, Yan X, Zhao H, Su L, Yang M, Gao C, Yao Y, Inoki K, Li D, Shao R, Wang S, Sahoo N, Kudo F, Eguchi T, Ruan B, and Xu H. Rapamycin directly activates lysosomal mucolipin TRP channels independent of mTOR. PLoS Biology 17: e3000252, 2019.

5. Bartolomé A, García-Aguilar A, Asahara S-I, Kido Y, Guillén C, Pajvani UB, and Benito M. MTORC1 Regulates both General Autophagy and Mitophagy Induction after Oxidative Phosphorylation Uncoupling. Molecular and Cellular Biology 37: e00441–00417, 2017.

6. Dai Y, Zheng K, Clark J, Swerdlow RH, Pulst SM, Sutton JP, Shinobu LA, and Simon DK. Rapamycin drives selection against a pathogenic heteroplasmic mitochondrial DNA mutation. Human Molecular Genetics 23: 637–647, 2014.

7. Gilkerson RW, De vries RLA, Lebot P, Wikstrom JD, Torgyekes E, Shirihai OS, Przedborski S, and Schon EA. Mitochondrial autophagy in cells with mtDNA mutations results from synergistic loss of transmembrane potential and mTORC1 inhibition. Human Molecular Genetics 21: 978–990, 2012.

8. Connolly BS, Feigenbaum ASJ, Robinson BH, Dipchand AI, Simon DK, and Tarnopolsky MA. MELAS syndrome, cardiomyopathy, rhabdomyolysis, and autism associated with the A3260G mitochondrial DNA mutation. Biochemical and Biophysical Research Communications 402: 443–447, 2010.

9. Flierl A, Reichmann H, and Seibel P. Pathophysiology of the MELAS 3243 transition mutation. Journal of Biological Chemistry 272: 27189–27196, 1997.

10. Sproule DM, and Kaufmann P. Mitochondrial encephalopathy, lactic acidosis, and strokelike episodes: Basic concepts, clinical phenotype, and therapeutic management of MELAS syndrome. Annals of the New York Academy of Sciences 1142: 133–158, 2008.

11. Barca E, Cooley V, Schoenaker R, Emmanuele V, DiMauro S, Cohen B, Karaa A, Vladutiu G, Haas R, Haas R, Van Hove J, Scaglia F, Parikh S, Bedoyan J, DeBrosse S, Gavrilova R, Saneto R, Enns G, Stacpoole P, Ganesh J, Larson A, Zolkipli-Cunningham Z, Falk M, Goldstein A, Tarnopolsky M, Camp K, Krotoski D, Engelstad K, Rosales X, Kriger J, Buchsbaum R, Thompson J, and Hirano M. Mitochondrial disease phenotypes of 999 patients in the North American Mitochondrial Disease Consortium (NAMDC) (P1.141). Neurology 90: P1.141, 2018.

12. El-Hattab AW, Adesina AM, Jones J, and Scaglia F. MELAS syndrome: Clinical manifestations, pathogenesis, and treatment options. Molecular Genetics and Metabolism 116: 4–12, 2015.

13. Berbel-Garcia A, Barbera-Farre JR, Porta Etessam JP, Martinez Salio A, Cabello A, Gutierrez-Rivas E, and Campos Y. Coenzyme Q 10 improves lactic acidosis, strokelike episodes, and epilepsy in a patient with MELAS (mitochondrial myopathy, encephalopathy, lactic acidosis, and strokelike episodes). Clinical Neuropharmacology 27: 187–191, 2004.

14. Paik K, Lines MA, Chakraborty P, Khangura SD, Latocki M, Al-Hertani W, Brunel-Guitton C, Khan A, Penny B, Rockman-Greenberg C, Rupar CA, Sondheimer N, Tarnopolsky M, Tingley K, Coyle D, Dyack S, Feigenbaum A, Geraghty MT, Gillis J, Van Karnebeek CDM, Kronick JB, Little J, Potter M, Siriwardena K, Sparkes R, Turner LA, Wilson K, Buhas D, and Potter BK. Health Care for Mitochondrial Disorders in Canada: A Survey of Physicians. Canadian Journal of Neurological Sciences 46: 717–726, 2019.

15. Rodriguez MC, MacDonald JR, Mahoney DJ, Parise G, Beal MF, and Tarnopolsky MA. Beneficial effects of creatine, CoQ10, and lipoic acid in mitochondrial disorders. Muscle and Nerve 35: 235–242, 2007.

16. Steriade C, Andrade DM, Faghfoury H, Tarnopolsky MA, and Tai P. Mitochondrial encephalopathy with lactic acidosis and stroke-like episodes (MELAS) may respond to adjunctive ketogenic diet. Pediatric Neurology 50: 498–502, 2014.

17. Johnson SC, Martinez F, Bitto A, Gonzalez B, Tazaerslan C, Cohen C, Delaval L, Timsit J, Knebelmann B, Terzi F, Mahal T, Zhu Y, Morgan PG, Sedensky MM, Kaeberlein M, Legendre C, Suh Y, and Canaud G. mTOR inhibitors may benefit kidney transplant recipients with mitochondrial diseases. Kidney International 95: 455–466, 2019.

18. McAlary L, Plotkin SS, and Cashman NR. Emerging Developments in Targeting Proteotoxicity in Neurodegenerative Diseases. CNS Drugs 33: 883–904, 2019.

19. Demers-Lamarche J, Guillebaud G, Tlili M, Todkar K, Bélanger N, Grondin M, P’Nguyen A, Michel J, and Germain M. Loss of mitochondrial function impairs Lysosomes*. Journal of Biological Chemistry 291: 10263–10276, 2016.

20. Joseph AM, Rungi AA, Robinson BH, and Hood DA. Compensatory responses of protein import and transcription factor expression in mitochondrial DNA defects. American Journal of Physiology - Cell Physiology 286: C867–875, 2004.

21. Menzies KJ, Robinson BH, and Hood DA. Effect of thyroid hormone on mitochondrial properties and oxidative stress in cells from patients with mtDNA defects. American Journal of Physiology - Cell Physiology 296: C355–362, 2009.

22. Iqbal S, and Hood DA. Oxidative stress-induced mitochondrial fragmentation and movement in skeletal muscle myoblasts. American Journal of Physiology - Cell Physiology 306: C1176–1183, 2014.

23. Garrido-Maraver J, Paz MV, Cordero MD, Bautista-Lorite J, Oropesa-Ávila M, de la Mata M, Pavón AD, de Lavera I, Alcocer-Gómez E, Galán F, Ybot González P, Cotán D, Jackson S, and Sánchez-Alcázar JA. Critical role of AMP-activated protein kinase in the balance between mitophagy and mitochondrial biogenesis in MELAS disease. Biochimica et Biophysica Acta - Molecular Basis of Disease 1852: 2535–2553, 2015.

24. Cotán D, Cordero MD, Garrido-Maraver J, Oropesa-Avila M, Rodríguez-Hernández Á, Izquierdo LG, Mata MDl, Miguel MD, Lorite JB, Infante ER, Jackson S, Navas P, and Sánchez-Alcázar JA. Secondary coenzyme Q 10 deficiency triggers mitochondria degradation by mitophagy in MELAS fibroblasts. The FASEB Journal 25: 2669–2687, 2011.

25. Morán M, Delmiro A, Blázquez A, Ugalde C, Arenas J, and Martín MA. Bulk autophagy, but not mitophagy, is increased in cellular model of mitochondrial disease. Biochimica et Biophysica Acta - Molecular Basis of Disease 1842: 1059–1070, 2014.

26. James AM, Wei YH, Pang CY, and Murphy MP. Altered mitochondrial function in fibroblasts containing MELAS or MERRF mitochondrial DNA mutations. Biochemical Journal 318: 401–407, 1996.

27. Tokuyama T, Hirai A, Shiiba I, Ito N, Matsuno K, Takeda K, Saito K, Mii K, Matsushita N, Fukuda T, Inatome R, and Yanagi S. Mitochondrial dynamics regulation in skin fibroblasts from mitochondrial disease patients. Biomolecules 10: 0, 2020.

28. Suárez-Rivero J, Villanueva-Paz M, de la Cruz-Ojeda P, de la Mata M, Cotán D, Oropesa-Ávila M, de Lavera I, Álvarez-Córdoba M, Luzón-Hidalgo R, and Sánchez-Alcázar J. Mitochondrial Dynamics in Mitochondrial Diseases. Diseases 5: 1, 2016.

29. Villa-Cuesta E, Holmbeck MA, and Rand DM. Rapamycin increases mitochondrial efficiency by mtDNAdependent reprogramming of mitochondrial metabolism in Drosophila. Journal of Cell Science 127: 2282–2290, 2014.

30. Shin YJ, Cho DY, Chung TY, Han SB, Hyon JY, and Wee WR. Rapamycin reduces reactive oxygen species in cultured human corneal endothelial cells. Current Eye Research 36: 1116–1122, 2011.

